# Sensitive Hormone and Neurotransmitter Detection with Carbon Flower Electrodes

**DOI:** 10.1101/2025.10.20.683291

**Authors:** Ines C. Weber, Diego U. Patino, Kuang-Jung Hsu, Yann Zosso, Adrian L.M. Düsselberg, Tianyang Chen, Kostas Parkatzidis, Shiyuan Wei, Alam Mahmud, Grégoire M.G.B.H. Bastide, Anna L. Remund, Mengfei Ashley Wu, Zhenan Bao

## Abstract

Carbon-based electrochemical sensors have attracted substantial attention for continuous health monitoring due to their high surface area, wide potential window, capability for repeated measurements, and compatibility with soft wearable electronics. However, they face challenges when detecting target biomarker concentrations in the low nanomolar range (*sensitivity*) and differentiating them in a mixture (*selectivity*), limiting their applicability in real scenarios.

Herein, we present sensitive and selective ‘carbon flower’ sensors, fabricated via a facile and patternable spray-coating process on soft, stretchable substrates. The carbon flowers—obtained through the synthesis of polyacrylonitrile and subsequent heat treatments—exhibit unique hierarchical morphologies, high surface area, and excellent conductivity, ideal for mass transport and electrochemical detection. We demonstrate the detection of estradiol, serotonin, melatonin, dopamine, uric acid, and ascorbic acid with detection limits as low as sub-nanomolar. The carbon flower sensors exhibit good repeatability across 100 cycles and over several weeks, robustness to pH and salt variations, and excellent performance in complex bio-fluids such as artificial saliva. In mixtures containing up to four analytes, they differentiate individual molecules, demonstrating the selectivity of the carbon flowers. This combination of sensitivity, selectivity, and mechanical compatibility makes carbon flower sensors well-suited for biomolecular sensing in soft, skin-conformable wearable electronic patches.

## Introduction

Following the success of the commercial continuous glucose monitoring devices, chemical sensors for continuous health monitoring have attracted substantial interest throughout the last decades [1–3]. Of particular interest is monitoring hormones and neurotransmitters due to the crucial roles they play in our brain, bi-directionally regulating body and behavior. For example, monitoring estradiol (E2), the primary form of estrogen, could provide valuable insights into bone [4] and muscle health [5], while it is relevant in female health for fertility assessment [6], hormone replacement therapy [4], or endometriosis/adenomyosis [7]. Measuring melatonin (Mel)—known as the sleep regulating hormone—could provide insights into body circadian rhythm and help manage sleep disorders [8]. Serotonin (5-HT) dysregulation has been associated with neurodevelopmental [9] disorders, gastrointestinal disorders [10], blood pressure regulation [11], kidney disease [11], and depression. Dopamine (DA) is important in reward-learning, motivation, and motor control [12], while its dysregulation has been associated with Parkinson’s, schizophrenia [13], attention deficit [14], and addiction [15]. Therefore, there is a clear need for regular and individualized assessment of these biomarker concentrations to effectively track health, adjust therapies, and facilitate diagnoses.

Currently, measuring these biomarkers involves invasive blood draws and subsequent analysis using expensive and bulky instruments such as liquid chromatography-mass spectrometry (LC-MS). These tests are infrequent and only done in case of strong clinical suspicion of disease. In contrast, low-cost portable and wearable chemical sensors offer a promising alternative for monitoring biomarker levels non-invasively, for example, through sweat, saliva, or interstitial skin fluid monitoring. However, the sensor requirements are extremely challenging for continuous sensing. Sensors need to be sensitive to detect low nanomolar or picomolar concentrations [16]; selective to distinguish between multiple analytes present at the same time; and stable to allow for repeated use in a medium without conditioning. At the same time, they should be stretchable to conform with the skin for more comfort, ideally low-cost, and be fabricated in a simple way.

Carbon-based electrochemical sensors have emerged as a promising material. Their operation principle relies on the oxidation and reduction of target molecules that result in measurable currents proportional to the analyte concentration [17]. This process is repeatable without the need to regenerate the surface, providing a significant benefit over biosensors that rely on bio-recognition elements and often face issues with stability and difficulty in regeneration due to strong binding affinity of the receptor to the target analyte, restricting the de-binding process [18]. Furthermore, carbon-based materials can exhibit high surface area and conductivity, providing abundant electroactive sites and conductivity ideal for sensing, while in some cases they have even been integrated into stretchable and wearable devices for on-skin and implantable applications [19]. Because of these significant advantages, carbon-based sensors have been studied by altering their morphology (e.g., carbon nanotubes, graphene, graphene oxide), combining them with metals and metal oxides, or enhancing performance through polymers (**Table S1**). However, these materials exhibit several critical limitations that hinder practical application. First, they demonstrate insufficient sensitivity, with reported 5-HT limits of detection (LODs) typically exceeding 10 nM while dermal sensing applications require 1 nM sensitivity [20]. Second, fabrication often involves complex synthetic procedures that impede scalability and require costly cleanroom processing for device integration. Third, many carbon-based materials suffer from batch-to-batch variability and limited control over their carbonization process. Finally, comprehensive selectivity assessment across multiple biomarkers is rarely conducted, limiting understanding of potential cross-reactivity in biological matrices.

Herein, we report a new class of carbon material, namely nanostructured carbon flowers (CFs), to address these challenges. CF are hierarchical carbon particles (∼1 µm diameter) obtained through a controllable, gram-scale protocol, where polyacrylonitrile (PAN) polymeric flower precursors are stabilized through sidechain cyclization and subsequent carbonization steps to form graphitic structures maintaining the hierarchical structures [21]. This process yields flower-shaped structures with readily accessible surfaces for electrochemical redox reactions and good electrical conductivity, previously demonstrated in electrocatalysis [22], pressure sensors [23], methane storage [24], and batteries [25]. The uniform particle size and homogeneous morphology enable robust and reproducible sensing performance, while also allowing straightforward patterning by spray-coating onto stretchable polystyrene-block-poly(ethylene-ran-butylene)-block-polystyrene (SEBS) substrates, thereby facilitating scalable and high-throughput fabrication of flexible sensors. We demonstrate that CF sensors exhibit capacitive performance and redox reversible character, enabling repeated, sensitive, and selective detection of multiple biomarkers including E2, 5-HT, Mel, DA, ascorbic acid (AA), and uric acid (UA), as well as mixtures thereof. The sensors demonstrate high stability, robustness to acid conditions and salt concentrations in the physiological range, ultra-low detection limits, and inherent selectivity across diverse analytes.

## Experimental

### Carbon flower synthesis

The CFs were synthesized following a procedure previously reported by our group [26]. In brief, acrylonitrile (≥ 99%, Sigma Aldrich, purified by percolation through basic alumina) and acetone were mixed in a 1:1 volume ratio with 1 mg azobisisobutyronitrile (≥ 99%, Sigma Aldrich) per mL of monomer. The solution was heated to 70 °C for 4 h without stirring in a sealed vial. The resultant PAN flowers were collected and washed with methanol and dried overnight under vacuum at 80 °C to remove any unreacted reaction components. Oxidative stabilization was done in air at 230 °C for 4 h to ensure the flowers retain their morphology during carbonization. Thereafter, the stabilized flowers were ramped at 2 °C/min to carbonization temperatures of 1000 (referred to as CF1000), 1250 (CF1250), and 1500 °C (CF1500) for 2 h in a tube furnace under nitrogen flow (100 mL/min). Carbon spheres were prepared from identical starting materials but stabilized under a nitrogen atmosphere followed by identical carbonization conditions at 1000 °C [22].

### Material characterization

An FEI Magellan 400 XHR Scanning Electron Microscope was used for scanning electron microscopy (SEM) with a current of 100 pA and a voltage of 5 kV. X-Ray diffraction was carried out with a Bruker D8 instrument and a Cu Kα source. Nitrogen sorption experiments were performed using an Autosorb iQ2 (Quantachrome) low-pressure gas sorption analyzer with 99.999% N_2_ at 77.35 K. Raman spectroscopy was run on a Horiba XploRA+ Confocal Raman System using a laser wavelength of 532 nm and a grating of 1200 gr/mm. The powders were pressed onto drop-cast SEBS substrates using a heat press to ensure flat films. For X-ray Photoelectron Spectroscopy (XPS, PHI VersaProbe 3), samples were electrically grounded to the sample stage inside the ultra-high vacuum chamber. Measurements were carried out using the monochromated Kα line of an aluminum X-ray source (1486.6 eV) with the analyser set at a pass energy of 224 eV for the survey spectrum and 112 eV for the high-resolution C1s and N1s. Bulk elemental analysis was carried out with a Thermo FlashSmart Soil NC Elemental Analyzer (EA), using ∼ 1 mg per sample. For optical images, a microscope (Leica Microsystems) was used.

### Device fabrication

#### Carbon flower ink

The CF ink was prepared by dispersing 45 mg of the respective CF material and 5 mg polyvinylidene fluoride (PVDF, *M*_w_ = 534,000 Da, Sigma Aldrich) in 100 mL chloroform. The mixture was initially sonicated overnight at room temperature, followed by probe ultrasonication for 20 min at 40% amplitude using a 2 s on/off pulse cycle (Cole-Parmer CP750 ultrasonicator, Vernon Hills, IL, USA). To remove large agglomerates, the dispersion was centrifuged at 1000 rpm for 1 min using a Sorvall LYNX 4000 centrifuge (Thermo Fisher Scientific, Waltham, MA, USA). The supernatant containing well-dispersed CFs was collected and used for subsequent electrode fabrication. The PVDF binder served to stabilize the CF dispersion and prevent particle settling during the spray coating process.

#### Sensor fabrication

The step-by-step fabrication protocol for CF sensors is illustrated in **Fig. S1**. This fabrication approach was designed to be straightforward and rapid, requiring no cleanroom facilities or expensive equipment. SEBS elastomer (Asahi Kasei Tuftec H1052, Tokyo, Japan) was dissolved in cyclohexane at 150 mg/mL and drop-cast onto glass slides, then dried overnight at room temperature to form a flexible substrate. Simultaneously, a Kapton tape mask containing eight electrode patterns (10.5 × 0.5 mm each) was laser-cut using an Epilog Fusion M2 CO_2_ laser cutter (Golden, CO, USA). The mask was applied to the dried SEBS film, and stretchable silver paste (ACI Materials SS1109, San Jose, CA, USA) was deposited using a blade-coating technique. The assembly was cured at 100 °C for 2 h, after which the Kapton mask was removed to reveal the patterned silver electrodes. A precision metal mask (15 mil Kovar alloy, Metal Etch Services, San Marcos, CA, USA) featuring eight electrode openings with alignment indicators was then positioned to overlap the silver electrodes while extending 2.5 mm beyond them onto the bare SEBS substrate. A magnet on the other side of the microscope slide helped to stabilize the metal mask. This configuration ensured CF deposition onto both the conductive silver regions to establish good electrical contact between the silver current collectors and CF active material, and the surrounding SEBS area, which serves as the sensing region. The CF ink (7 mL) was spray-coated at 80 °C from approximately 10 cm height using an airbrush system (Master Airbrush Model SB84 with 0.3 mm tip). Following spray coating, the metal mask was removed, and the non-active areas were encapsulated either by spin-coating an additional SEBS layer (80 mg/mL in toluene) for complete device sealing or, for rapid prototyping, by applying Kapton tape. Electrical connections were established using conductive z-axis tape to interface the CF electrodes with a flat cable connector.

### Sensing tests

Electrochemical measurements were performed using a CH Instruments 760E potentiostat equipped with a CHI200 picoamp booster and Faraday cage to minimize electrical interference. A conventional three-electrode configuration was employed, comprising an Ag/AgCl (3 M KCl) reference electrode, platinum wire counter electrode, and CF working electrode. All measurements were conducted in nitrogen-degassed 1x phosphate-buffered saline (PBS, pH 7.2) at room temperature. The chemicals 17β-estradiol (> 97%, TCI Chemicals) and melatonin were prepared in ethanol, while serotonin hydrochloride (≥ 98%, Sigma Aldrich), L-ascorbic acid (≥ 98%, Sigma Aldrich), uric acid (98%, AmBeed), and dopamine hydrochloride (≥ 98%, Sigma Aldrich) were dissolved in PBS buffer. All analyte solutions were prepared immediately before measurements and added sequentially to the nitrogen-degassed PBS buffer. Gentle mixing was achieved by brief nitrogen purging. A standardized wait time of 60 s was implemented between consecutive square wave voltammetry (SWV) scans, except during repetitive cycling experiments where a 5-min interval was employed to assess electrode stability and reproducibility. Measurements in physiological conditions were conducted using artificial saliva (Pickering Laboratories 1700-0304, pH 6.8, containing <0.5% potassium chloride). Sensors were rinsed with DI water prior to measurements.

The following measurement parameters were employed for cyclic voltammetry (CV), electrochemical impedance spectroscopy (EIS), and SWV:

- CV: Measurements were performed over a potential window of -0.4 to +1.3 V vs. Ag/AgCl at a scan rate of 0.1 – 2 V/s with a sampling interval of 0.001 V. For electrochemically active surface area determination, DA (20 µM in PBS) was employed as a redox probe molecule. After use, sensors were stored in a nitrogen box.
- EIS: Measurements were conducted at the open circuit potential over a frequency range of 100 kHz to 0.1 Hz with an alternating current amplitude of 10 mV.
- SWV: SWV was performed with a pulse amplitude of 0.025 V, frequency of 5 Hz, and potential increment of 0.004 V in the range of 0 – 0.8 V. For analytes testing, SWV was selected due to its advantages over other voltametric methods, including high sensitivity, reduced capacitive current, and rapid acquisition times

#### Comparative Electrode Materials

For performance benchmarking, carbon sphere electrodes were fabricated using identical procedures as the CF sensors to enable direct morphology-dependent comparisons. Additionally, glassy carbon electrodes (GCE, 2 mm diameter, CH Instruments Inc., Austin, TX, USA) served as conventional carbon electrode controls. Prior to use, GCE surfaces were mechanically polished using a sequential alumina powder treatment (0.3 μm followed by 0.05 μm particle sizes) from the electrode polishing kit (CH Instruments Inc.). Following polishing, electrodes were thoroughly rinsed with deionized water.

Acidity and salt tests: the pH was adjusted by adding stepwise 10 μL of 0.1M HCl into 10 mL PBS containing 500 nM E2. The actual pH values were determined with a pH meter. For salt testing, 5-50 mM of KCl were added to the PBS solution containing 500 nM E2.

### Data analysis

For electrochemical SWV data analysis, background subtraction was performed using the response obtained in blank PBS electrolyte under identical conditions. Following background correction, a linear baseline was established between the start and end points of the current peak and subtracted to isolate the analyte signal. Data smoothing was applied over 20 measurement points using the ‘movmean’ function on MATLAB. The analytical sensitivity was defined as the slope of the peak current as a function of concentration. The limit of detection (LOD) was calculated as LOD = 3 σ/S, where σ represents the standard deviation of the baseline noise and S is the analytical sensitivity.

The electrochemically active surface area (ECSA) was determined using the Randles-Ševčík equation applied to DA cyclic voltammetry data collected at scan rates ranging from 10 to 100 mV/s. The diffusion coefficient of DA (7.6 × 10^-6^ cm^2^/s) and the peak current versus the square root of scan rate were used to calculate the effective electrode area. Additionally, the slope of log(peak current) versus log(scan rate) was analyzed to distinguish between diffusion-controlled (slope ≈ 0.5) and adsorption-controlled (slope ≈ 1.0) electrochemical processes. Calibrated sensor response corresponds to the sensor current compared to the single analyte measurement on that same sensor. Data analysis was performed using MATLAB (2023). Number of replicates is shown with individual dots (for less than 6 replicas) or as error bars (for more than 6 replicas).

## Results and Discussions

The rationale behind the development of CF sensors is to present a novel, high-performance carbon sensing material alongside the facile fabrication of multi-electrode, flexible sensors. Our sensor fabrication employs a straightforward spray coating technique to apply CFs with unique morphology, high conductivity, and high surface area onto stretchable substrates, as shown in **Fig. 1a,b**. The metal mask enables efficient patterning and multi-electrode sensor arrays. The simplicity of the spray-coating technique enables the fast deposition of a multitude of sensors onto a substrate of choice, here a stretchable SEBS substrate (**Fig. 1c**). This flexibility is essential for health monitoring applications, where sensors must conform to skin movement when placed directly on the body, as well as for implantable devices such as CGM needles that require mechanical compliance with tissue.

**Fig. 1.**
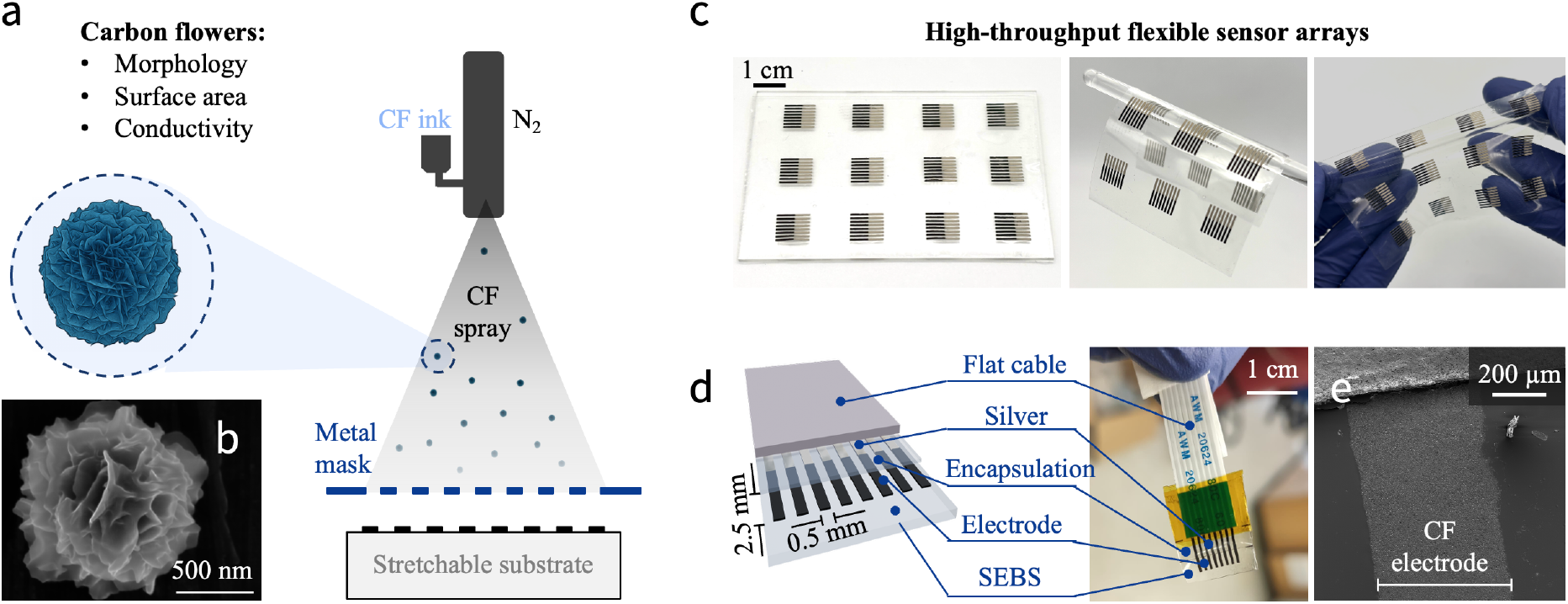
Carbon flower sensors: concept, design, properties. **(a)** Schematic of spray-coated carbon flower sensors with the capability of biomolcculc recognition for wearable, skm-conformablc sensors. **(b)** Representative SEM image of a carbon flower microparticle.**(c)** Photograph of high-throughput flexible and stretchable sensor arrays. **(d)** Schematic and photograph of the sensor, (e) SEM image of the working electrode.

A schematic illustration of the device architecture, together with a photograph, is depicted in **Fig. 1d**. Each sensor comprises eight electrodes with exposed surface area of 2.5 × 0.5 mm. Stencil-printed stretchable silver interconnects connect the CF sensors to a flat cable for signal read-out. The resulting electrodes exhibit a homogeneous surface with well-defined geometric dimensions (**Fig. 1e**). Further insights into electrode sizing, including top-view optical images and SEM images of the sensor surface and cross-section are shown in **Fig. S2**.

Overall, this fabrication technique offers multiple advantages: it is cost-effective, consumes low power, reduces contamination risk, and simplifies fabrication steps. The sensing electrodes are based on the CF ink; the CF synthesis follows an established protocol, starting with the one-step synthesis of PAN flowers, followed by subsequent stabilization and carbonization [26]. This fabrication allows good control over the flower morphology and conductivity. Overall, the ease of this method, combined with the excellent sensitivity, selectivity, and stability performance of the CFs for electrochemical sensing, as shown below, are key merits of this work.

### Characterization of carbon flowers

The CF characterization (prior to formulating the electrode spray-coating ink) is presented in **Fig. 2**. SEM imaging reveals flower-like particles comprised of ‘petals’ (**Fig. 2a-c**). The particles exhibit a distinctive flower-like morphology with excellent uniformity in size, showing average diameters of 0.88 ± 0.08 µm, 0.86 ± 0.06 µm, and 0.80 ± 0.07 µm for the CF1000, CF1250, and CF1500, respectively (histograms in **Fig. S3**). The size reduction is consistent with previous works on carbonizing PAN fibers [27]. No visible morphological changes occur across carbonization temperatures, as expected given the use of identical polymeric and stabilized precursors. The uniform particle size distribution and consistent morphology are crucial to ensure consistent electrode performance and sensor-to-sensor reproducibility.

**Fig. 2.**
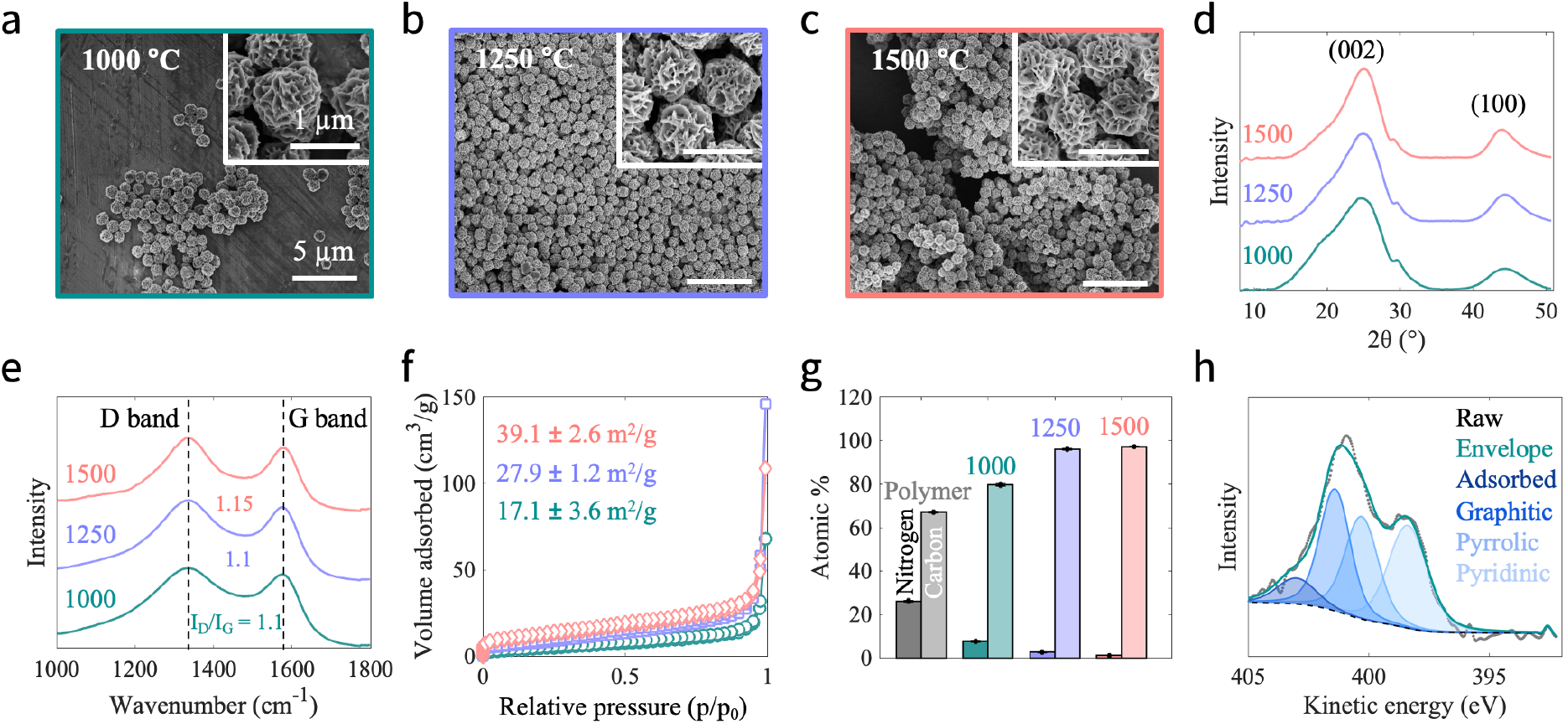
Morphology, composition, and structure of carbon flowers. **(a-c)** SEM images of the carbon flowers carbonized at 1000 °C, 1250 °C, and 1500 °C. **(d)** XRD patterns. **(e)** Raman spectra. **(f)** Nitrogen adsorption isotherms and BET surface area (three replicas). **(g)** Elemental composition (wt% C and N) of polymeric flowers and carbon flowers. Error bars and measurement points show three replicas. **(h)** XPS high-resolution nitrogen spectra of1000 °C carbon flowers. From left to right: adsorbed, graphitic, pyrrolic, pyridinic nitrogen.

The X-ray diffraction patterns (**Fig. 2d**) reveal broad diffraction peaks centered around ∼25° and ∼42°, corresponding to the (002) and (100) reflections of the graphitic regions, consistent with prior studies on CFs [21]. With increasing carbonization temperature, the sharpness of the (002) peak slightly increased, indicating enhanced crystallinity and improved graphitic ordering [27], which directly correlates with higher electrical conductivity [29].

The Raman spectra (**Fig. 2e)** show prominent D and G peaks at wavenumbers of 1340 cm^-1^ and 1580 cm^-1^. The D/G intensity ratio > 1 indicates highly defective carbon structures, with minimal differences between different carbonization temperatures. The incorporation of a high defect density is generally beneficial for electrochemical sensing applications, as these serve as sensing active sites [30].

To evaluate the surface area and porosity of the CFs, nitrogen adsorption-desorption isotherms were measured at 77 K (**Fig. 2f**). All CF exhibited type-IV isotherms, consistent with existing literature [26], indicating the presence of micro and mesopores. The calculated surface area reached up to 39.1 m^2^/g and decreased with carbonization temperatures. While carbon materials with higher surface areas, usually requiring an activation process [31, 32], are frequently reported, the hierarchical petal structure of our CFs provides enhanced accessibility for biomolecules compared to conventional high-surface-area carbons that rely on micropores < 2 nm, which restrict the diffusion of larger biomarkers.

The bulk chemical composition of the flowers determined by elemental analysis (EA, **Fig. 2g**) shows that as-prepared polymeric flowers consist of 67.1 ± 0.2 wt% C and 26.1 ± 0.3 wt% N, which corresponds well with the theoretical C/N ratio of PAN. Upon carbonization, the carbon-to-nitrogen ratio systematically increases with temperature, reaching up to 97.1 wt% C and 1.1 wt% N in CF1500. This change in carbon to nitrogen ratio reflects the thermal degradation of nitrogen-containing functional groups at higher temperatures. The progressive increase in carbon content aligns with the enhanced crystallinity observed in XRD analysis, as both indicate improved graphitic ordering with temperature and higher conductivity, critical for efficient electron transfer in electrochemical sensing applications.

The surface chemical composition of the CF was further examined with X-ray photoelectron spectroscopy (XPS, **Fig. 2h** and **Fig. S4**). The survey scans and tabulated values confirm the EA results, showing increased carbon and decreased nitrogen content with temperature, in line with previous reports [22]. The total amount of nitrogen in the CF1000 sample was measured to be 5.55 at%, in good agreement with the bulk EA measurement (7.5 wt%). The high-resolution N1s spectrum shown in **Fig. 2h** reveals the presence of pyridinic (398.3 eV), pyrrolic (400.3 eV), graphitic (401.4 eV), and CO_2_ or H_2_O adsorbed nitrogen species (403 eV [33]). The survey scans indicate the existence of C, O, and N elements. While the N-doping has shown advantageous roles in electrocatalysis, higher carbonization temperatures result in a higher content of sp^2^ carbon in the resulting CFs (Fig. S4), hence leading to higher conductivity, also advantageous for sensing.

In summary, the CFs possess several key characteristics that make them promising for electrochemical sensing applications: (i) uniform flower-shaped morphology with open petal structures that facilitate biomolecule diffusion and accessibility, (ii) good crystallinity and high sp^2^ carbon content for efficient electron transfer, and (iii) high defect density providing abundant active sites. These combined properties position the CFs as excellent electrode materials for sensitive and selective detection of biological analytes, as demonstrated in the following chapters.

### Electrochemical properties

The electrochemical performance of CF sensors is shown in **Fig. 3**. Cyclic voltammetry was used to characterize the electrochemical behavior (**Fig. 3a**). In blank PBS, no discernible redox peaks are observed. The rectangular shape of the I-V curves over a wide potential range confirms excellent capacitive behavior of the electrodes, needed for sensitive measurements of Faradaic processes. The symmetrical linear increase in anodic and cathodic currents at 0.5 V indicates a predominance of capacitive processes (R^2^ > 0.99, **Fig. S5a**) and rapid and reversible charging and discharging of the electric double layer. The low standard deviation across eight electrodes demonstrates negligible inter-electrode variability. Notably, higher carbonization temperatures lead to increased capacitance and larger currents while all the electrodes are kept at similar thickness of 18 µm (**Fig. 3a** inset, **Fig. S5b**,**c** for full scans). This improvement is consistent with the observed increase of surface area via BET analysis as well as the higher crystallinity and hence conductivity observed in XRD, EA, and XPS analysis. Similarly, the variation in electrode size affects capacitance: reducing the size of the electrode decreases the capacitance (**Fig. S6a**,**b**). Reversibility is confirmed with DA as a redox-active probe (**Fig. S5d**), showing a small oxidation and reduction peak separation of 60 mV. A linear correlation (R^2^ = 0.97) is observed between the logarithm of the scan rate and current, with a slope of 0.53, indicating a diffusion-controlled process (**Fig. S5e**). The electrochemical active surface area (ECSA), calculated using the Randles-Ševčík equation, is 3.5 mm^2^, exceeding the geometrical working electrode area (1.25 mm^2^), attributed to the hierarchical petal morphology of the CFs.

**Fig. 3.**
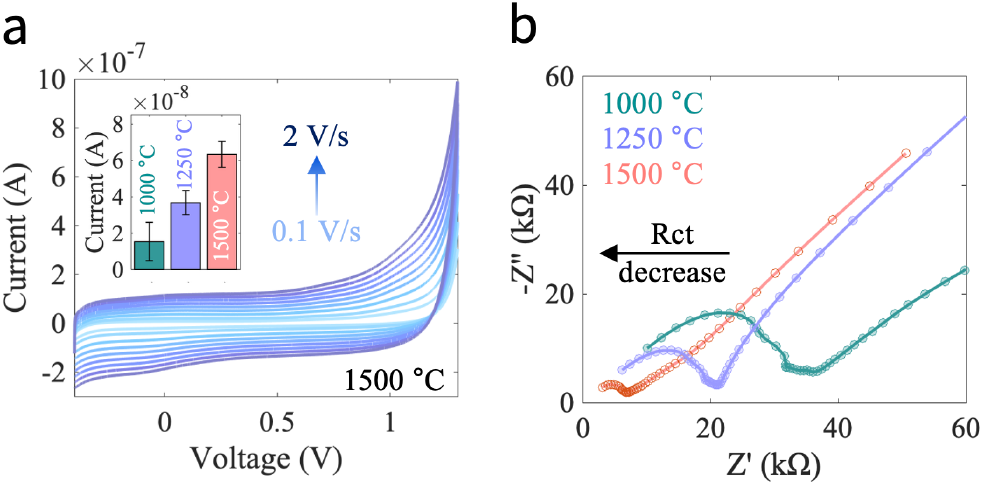
Electrochemical characterization. **(a)** Cyclic voltammograms of sensors fabricated from 1500 °C CF. The current at 0.5 V for 1 V/s scans, averaged over X electrodes, is shown in the inset. **(b)** Nyquist plot.

Electrochemical impedance spectroscopy (EIS) was employed to further elucidate the properties of the CF electrodes at the open circuit potential (**Fig. 3b**). The decrease in impedance observed in the Nyquist plots at elevated carbonization temperatures suggests enhanced electrochemical catalytic activity and electrode conductivity, which is in line with the observations obtained from Fig. 3a, as well as the larger sp^2^ carbon contributions in the CF1500. Similar trends were observed by reducing the electrode size, resulting in an increase in the charge transfer resistance (R_ct_) for smaller areas (**Fig. S6c**).

In summary, the electrochemical analysis is strongly correlated to the material characterization findings. The CF1500 exhibit superior performance, likely due to improved graphitic ordering and higher conductivity, which seems to out-weigh the presence of more nitrogen sites of the lower temperature CFs. The sheet resistance of spray-coated electrodes was measured to be 139.1, 34.3, and 4.2 kΩ for CF1000, CF1250, and CF1500, respectively (averaged over 8 electrodes). Owing to its beneficial properties, all subsequent sensing experiments utilized the CF1500 material.

### Estradiol detection: sensitivity, repeatability, robustness

Building on the promising characteristics of CF sensors, we exemplify their capability for detecting low biomarker concentrations through the analysis of E2 using SWV. As demonstrated in **Fig. 4a**, a pronounced peak is observed around 0.51 V for all concentrations ranging from 1 to 10 nM, exhibiting a linear relationship between peak current and concentrations (inset). Notably, the limit of detection reached 300 pM, which significantly outperforms recent findings in literature (**Table S1**). Reducing electrode size leads to a worse (i.e., higher) LOD (**Fig. S6d**), in line with the CV and EIS measurements in Fig. 3. This high sensitivity is substantially better compared to similarly prepared carbon spheres (**Fig. 4b**), which exhibit lower BET surface areas (9.7 m^2^/g) and lack the unique hierarchical morphology of our CFs, as shown in SEM and sensing data (**Fig. S7a**,**b**). Moreover, glassy carbon electrodes (GCE) demonstrate considerably lower sensitivity (**Fig. S7c**), further underscoring the superior electrochemical performance of the CF sensors.

**Fig. 4.**
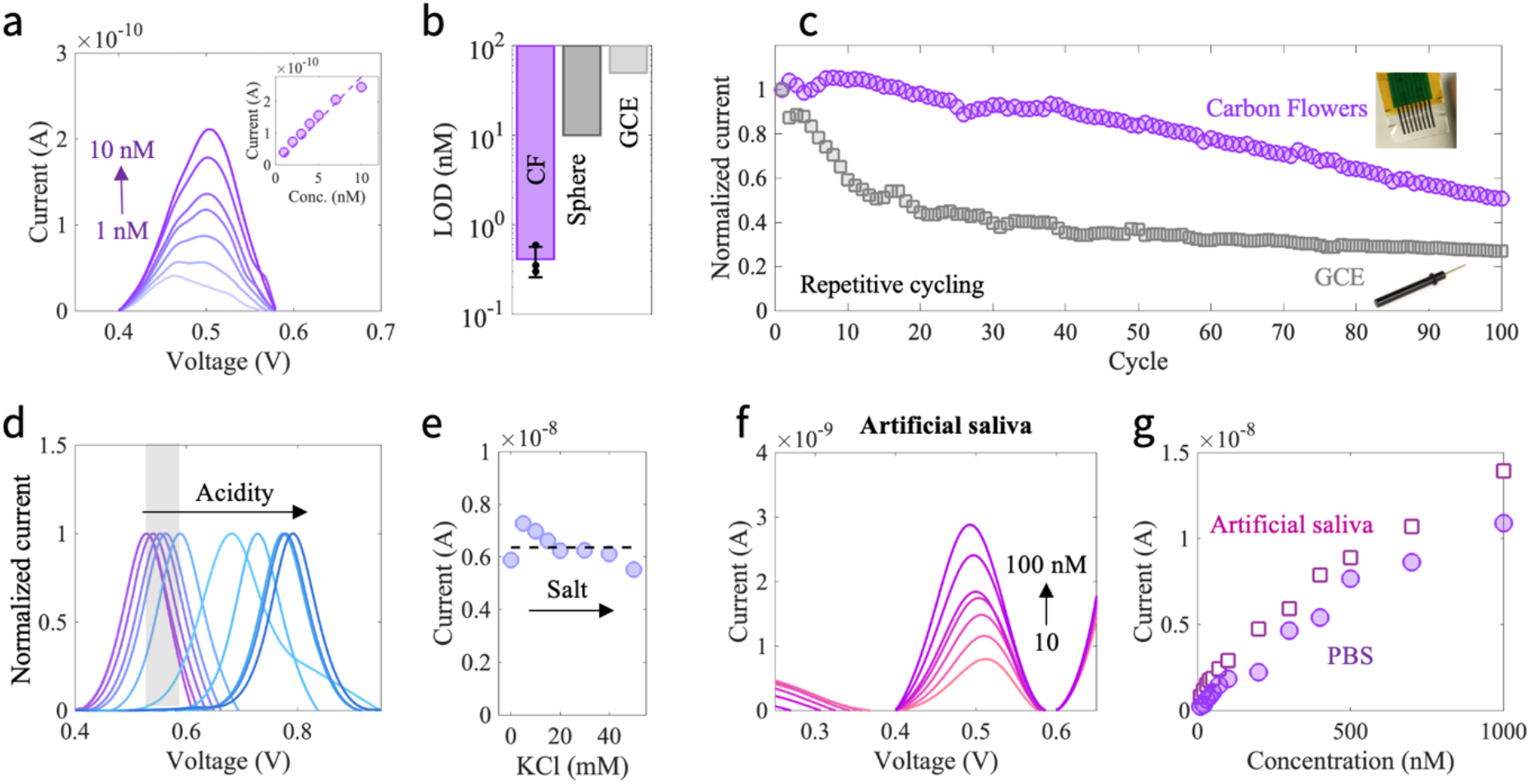
Performance characteristics for estradiol detection. **(a)** Detection of low estradiol concentrations with scatter plot (inset) and **(b)** limit of detection (LOD) in comparison to carbon spheres and GCE. **(c)** Normalized peak current response during repetitive cycling of CF and GCE electrodes. Normalized current response for 500 nM estradiol at varying pH **(d**, physiological range indicated in grey) and salt levels **(e). (f. g)** Estradiol detection in artificial saliva.

In addition to their exceptional sensitivity, CF sensors showed better stability during repetitive testing cycles in solution compared to GCE, even without stirring or refreshing the solution (**Fig. 4c**). While a gradual decrease in peak current is observed across the cycles, the CF sensors maintain substantial responsiveness, in contrast to the more rapid deterioration of GCE current peaks (see SWV full scans in **Fig. S8a**,**b**). This enhanced stability may be related to the hierarchical morphology of CFs compared to the flat GCE surface, though further investigation is needed.

To assess the utility of these sensors in physiological applications, we also investigated their performance under varying pH levels and salt concentrations, given the natural variability of bodily fluids. As shown in **Fig. 4d,e**, varying the pH from 7.2 to 2.9 resulted in a substantial anodic peak shift of 260 mV, accompanied by a notable increase in peak current of 70% (see raw data **Fig. S8c**). The potential shift is likely due to increased protonation of the phenolic groups, rendering oxidation less favourable, and is consistent with previous literature on DA sensing [34]. Remarkably, the sensors demonstrate minimal changes in peak position within the physiologically relevant pH range for saliva, indicated by the grey bar in Fig. 4d.

Potassium chloride was used to assess the impact of salt concentration on E2 detection (**Fig. 4e**; raw data **Fig. S8d**). KCl concentrations ranging from 0 – 50 mM were added to PBS buffer (baseline: 2.7 mM KCl), encompassing physiologically relevant salt levels. The E2 peak current remained stable at 6.4 ± 0.5 nA across this concentration range, demonstrating robust sensor performance under varying ionic conditions. Salt concentrations did not significantly affect the peak potential and, in some cases, slightly enhanced the peak current response.

Finally, we tested the sensor’s performance in artificial saliva with added 10 – 100 nM E2 (**Fig. 4f,g**). All tested concentrations are successfully detected, and the overall current response is higher in artificial saliva compared to PBS (**Fig. 4g** and **S9**). This may be attributed to both the higher salt levels as well as the more acidic conditions of artificial saliva (pH 6.8) compared to the pure PBS, in line with the pH-related observations outlined in Fig. 4d,e. Overall, these findings confirm that the CF sensors deliver reliable performance across various pH levels, salt concentrations, repetitive testing cycles, and in complex media such as artificial saliva.

### Carbon flowers for multiple biomarker detection

Having demonstrated the exceptional performance of CF sensors for E2 detection across various conditions, we next explored their versatility by expanding the analyte concentration range and investigating their capability for detecting multiple biomarkers of clinical relevance. To enable direct comparison with other analytes, E2 measurements were extended to concentrations up to 100 nM (**Fig. 5a**). The pronounced peak observed at 0.51 V maintained excellent linearity across this broader range, with maximum peak currents following a strong linear relationship with E2 concentration (R^2^ = 0.99).

**Fig. 5.**
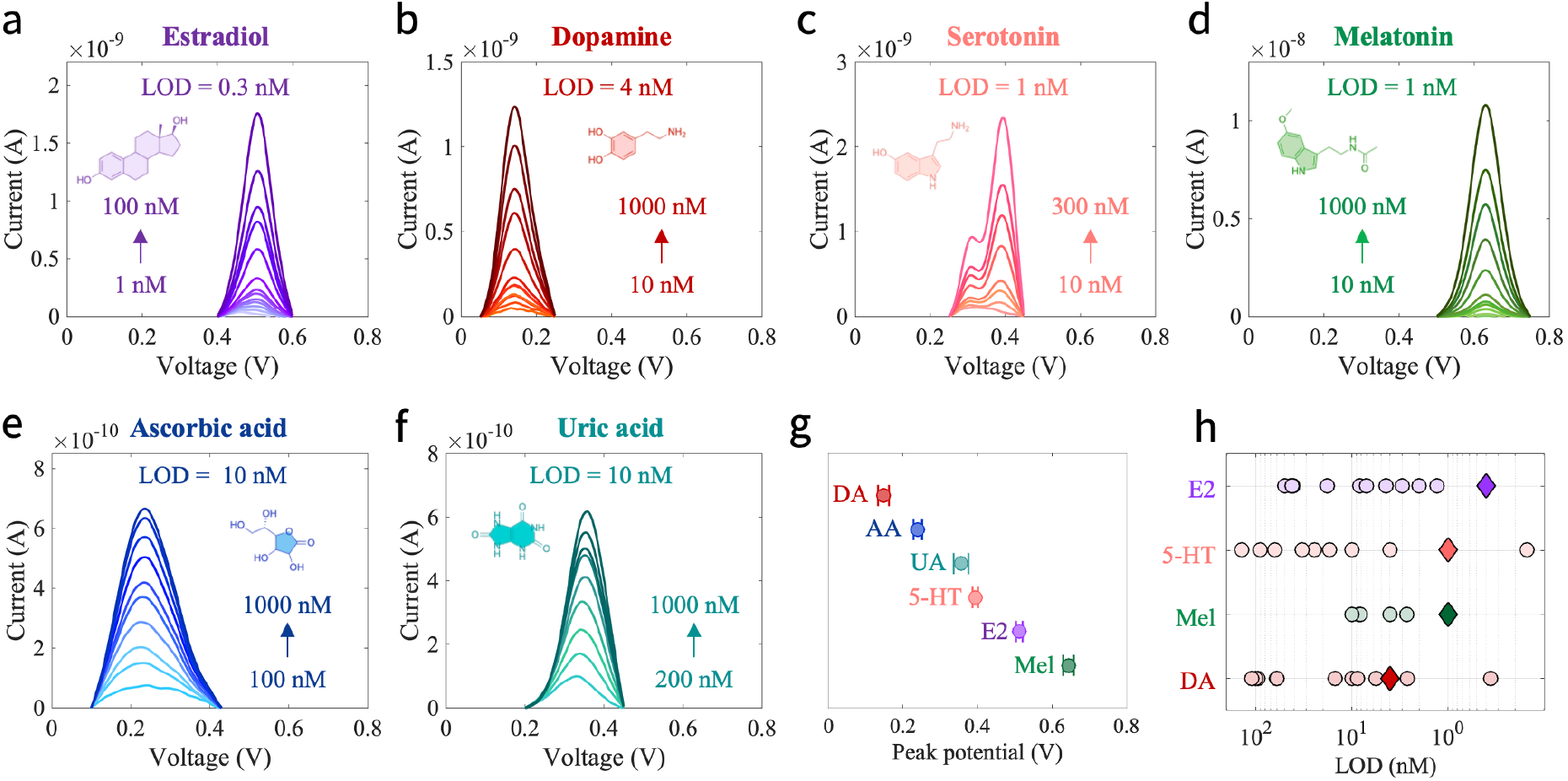
Detection of various biomarkers using carbon flower sensors via square wave voltammetry. Detection of estradiol (E2) **(a),** dopamine (DA) **(b),** serotonin (5-HT) **(c),** melatonin (Mel) **(d),** ascorbic acid (AA) **(e),** and uric acid (UA) **(f). (g)** Peak potentials at 1000 nM for each analyle. averaged over eight channels. Error bars represent standard deviation across all eight channels **(h)** LOD literature comparison of carbon-based electrochemical sensors since 2020 (representing Table SI). Diamonds indicate CFs.

Beyond E2, our study shows that CFs effectively detect various biomarkers (**Fig. 5b-f**) with high sensitivity. These include DA (peak at 0.14 V, LOD = 4 nM), 5-HT (0.33 and 0.38 V, LOD = 1 nM), and Mel (0.63 V, LOD = 1 nM), alongside AA (0.23 V, LOD = 10 nM), and UA (0.33 V, LOD = 10 nM). The raw data for all analytes is shown in **Fig. S10a-f**. The observed double-peak for 5-HT is in agreement with previous literature, where it has been attributed to oxidation first to carbocation, followed by a further oxidation to quinone imine [35]. Similar observations were made for Mel, showing additional, but much smaller, oxidation peaks that did however not affect mixture measurements (**Fig. S10g**). Importantly, all biomarkers show a linear response over the tested range and exhibit highly sensitive behavior (**Fig. S10h**). While UA and AA commonly occur in the µM concentration range, the former 4 analytes occur in the low nM or even pM range and require high sensitivity.

The distinct peak potentials, averaged over 8 electrodes, demonstrate a clear peak separation between most analytes and excellent reproducibility (**Fig. 5g**). Current responses at 1 μM concentration, averaged over all 8 electrodes, confirmed that CF sensors exhibited the highest response to E2 while maintaining consistent performance across the electrode array (**Fig. S10g**).

The detection range covers the physiological concentration range of most biomarkers in non-invasive body fluids, though lower detection limits remain needed for E2 and Mel in sweat and saliva. Importantly, the limits of detection are competitive and outperform most existing carbon-based electrochemical sensors (**Fig. 5h**, logarithmic scale, and **Table S1**), while maintaining the advantages of straightforward fabrication and preserved softness and stretchability for wearable applications.

### Multi-analyte detection: selectivity

In real applications such as saliva and sweat, these biomarkers co-occur; this co-occurrence causes significant challenges in detecting individual analytes in a mixture. Hence, we tested the CF sensors for mixtures of two up to four analytes. **Fig. 6a** shows the sensor’s performance when detecting varying concentrations of E2 (10 – 500 nM) in the presence of a fixed concentration of Mel (500 nM). Remarkably, despite orders of magnitude higher Mel concentrations, the sensor was still capable of detecting E2 in the low nanomolar range, as evidenced by the formation of a noticeable shoulder peak. In addition, the Mel peak exhibited only small reduction across repeated scans (from 8.8 to 3.1 nA) consistent with the repeatability observed for E2 in Fig. 5c.

**Fig. 6.**
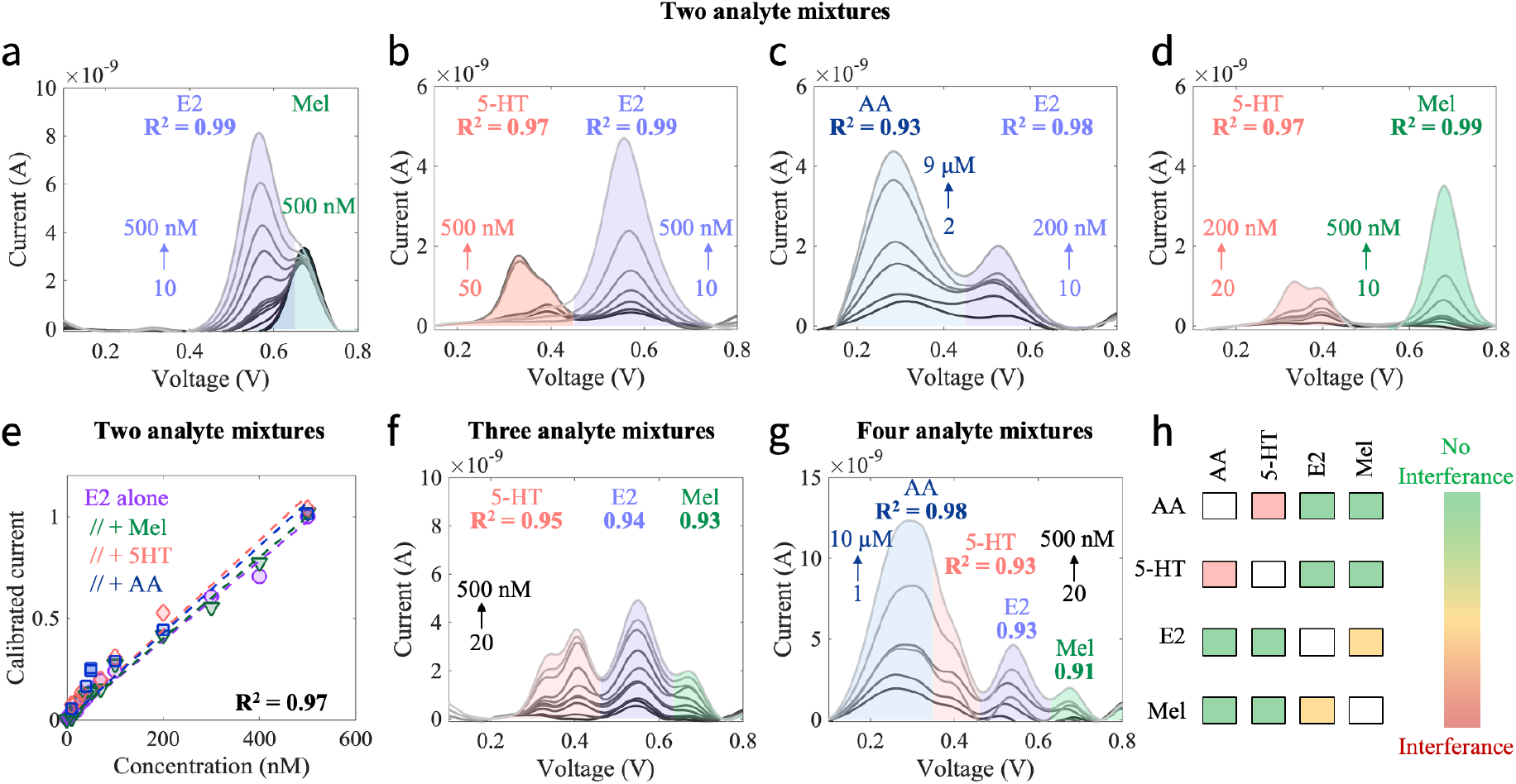
Selectivity. Estradiol detection in the presence of 500 nM melatonin **(a),** as well as varying serotonin **(b)** and ascorbic acid concentrations **(c). (d)** Simultaneous serotonin and melatonin measurement, **(e)** Peak current responses for two analyte mixtures normalized against single-analyte measures, **(f)** Three-analyte mixture of serotonin, estradiol, and melatonin (20-500 nM). **(g)** Four-analyte mixture of ascorbic acid (1-10 p.M). serotonin, estradiol, and melatonin (each 20 - 500 nM). **(h)** Interference summary.

To further challenge our sensors, we tested their capability to simultaneously detect E2 and 5-HT with both varying concentrations (**Fig. 6b**). Notably, the two analytes were clearly separated, achieving a high coefficient of determination greater than 0.97. Similar trends were observed with E2 in the presence of AA (**Fig. 6c**). Finally, the detection peaks for 5-HT and Mel remained completely separated (**Fig. 6d**).

Of note is that the measurements shown in Figs. 6b,c,d and 5c were conducted with the same sensor over a period of 17 days. The maintained current data of 4.63 ± 0.09 nA for 500 nM E2 demonstrates the sensor’s consistent performance despite the sensor storage in dry conditions. The absolute current may differ between individual sensors, as seen with higher currents in Fig. 6a. However, in real settings, the current response is typically normalized, as done here with the absolute response to E2 measurements from single-analyte measurements. The calibrated current response as a function of concentration depicted in **Fig. 6e** demonstrates excellent agreement between the measurements of E2 alone and in mixture with 5-HT, Mel, and AA, all exhibiting nearly identical sensitivity. This indicates that E2 detection is minimally impacted by the presence of other molecules, suggesting that the sensors do not suffer from competitive interactions over reactive sites, a critical factor in maintaining sensitivity and selectivity.

Finally, we tested the performance in three-analyte (**Fig. 6f**) and four-analyte (**Fig. 6g**) mixtures. Even in the three-analyte scenario, clear separation remained evident for 5-HT, E2, and Mel. In the case of four analytes, the 5-HT and AA peaks merge, complicating their differentiation. While the R^2^ values are lower in the four-analyte mixture, they remain relatively high, which however in case of AA and 5-HT is partially attributed to the steadily increasing concentrations.

These results, summarized in **Fig. 6h**, demonstrate the sensor’s effective analyte discrimination, particularly for E2 and Mel detection. The CF sensors exhibit promising inherent selectivity. We envision combining these sensors with other sensing materials in sensor arrays to further enhance the selectivity of electrochemical sensing devices for practical applications.

## Conclusions

CFs are effective, highly promising materials for electrochemical sensing. Through a simple fabrication technique of spray-coating, these materials can be employed as flexible sensors with high throughput and at low cost, without requiring controlled and expensive cleanroom facilities. The CFs exhibit unique morphology and a defect-rich structure. Higher carbonization temperatures lead to higher surface areas and higher carbon content, as evidenced by nitrogen adsorption, elemental analysis, and XPS. As sensors, the CFs exhibit excellent electrochemical stability and reversibility, with higher conductivity flowers at 1500 °C showing superior electrocatalytic activity. When tested with numerous analytes, the CFs demonstrate high sensitivity in the nanomolar range with excellent batch-to-batch and inter-electrode reproducibility and LODs exceeding currently reported carbon materials. CF sensors retain their sensing properties quite well with repetitive cycling and are also robust to varying pH and salt conditions, while they maintain their high performance in artificial saliva. Finally, they show inherent selectivity and can detect multiple analytes simultaneously in complex mixtures with up to four analytes, which is required for practical applications. Overall, CFs show great potential to be employed as an active sensing material by the wider community for numerous biomedical applications.

## Acknowledgements

This project was supported by the Stanford Wearable Electronics Initiative (eWEAR) seed grant and the Swiss National Science Foundation (SNSF) through postdoctoral fellowships (I.C.W: P500PT_214498; K-J. H: P500ON_230529; K.P: P500PN_222266). Part of this work was performed at the Stanford Nano Shared Facilities (SNSF), supported by the National Science Foundation under award ECCS-2026822. Additional support was provided by the U.S. Department of Energy, Office of Science, Office of Basic Energy Sciences, Chemical Sciences, Geosciences, and Biosciences Division, Catalysis Science Program, through the SUNCAT Center for Interface Science and Catalysis (D.U.P). We thank A. Chen and A.G. Zhang for technical assistance on data collection, C. Xu for scientific advice, and E. Luis for feedback on the paper.

## Data availability

The data that support the findings of this study are available from the corresponding author upon request.

