## Supplementary Information for "Sensitive Hormone and Neurotransmitter Detection with Carbon Flower Electrodes"

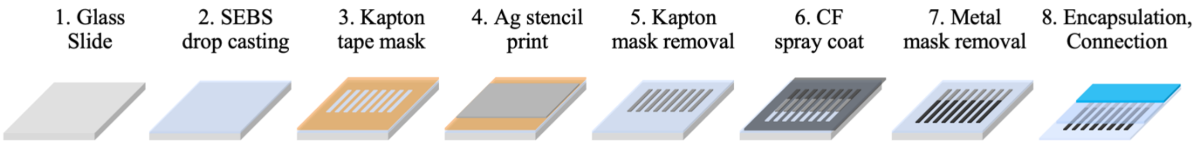


**Fig. S1| Sensor fabrication.** A glass slide is used as substrate (1) for SEBS drop-casting (2). A Kapton tape mask (3) defines the pattern for the stretchable Ag stencil-printing (4, 5). The CF ink is spray-coated using a metal mask (6). Encapsulation with spin-coated SEBS (7) and connection through z-axis tape and a flat cable conclude the fabrication (8).

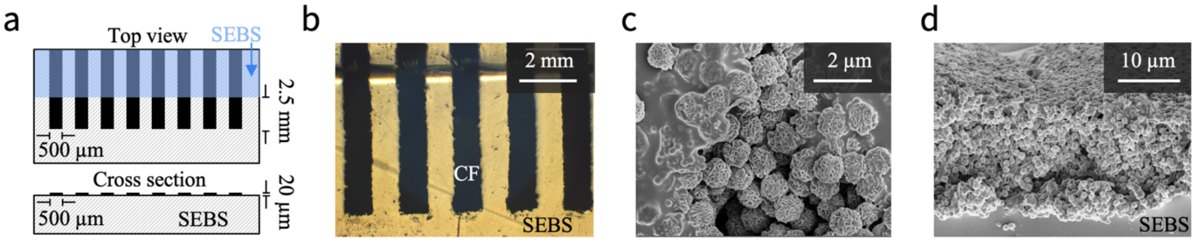


**Fig. S2| Sensor design and characterization. (a)** Schematic top and cross section view of the sensor layout. **(b)** Optical microscope image of the fabricated sensor (top view). **(c)** SEM image of sensor top view. **(d)** Cross-sectional SEM image of the sensor structure.

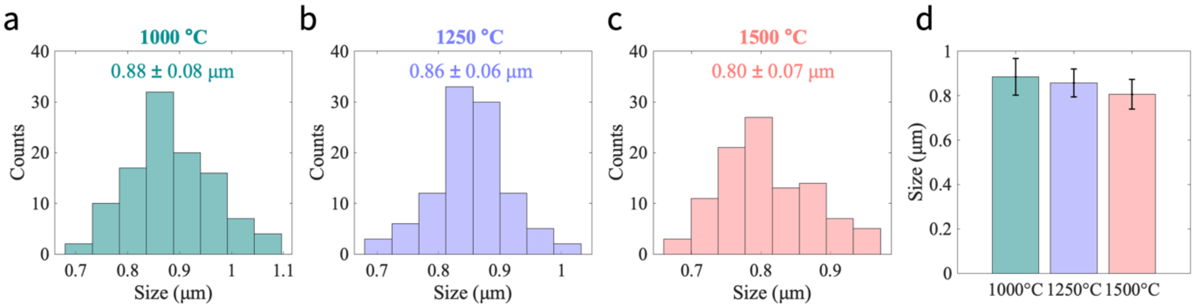


**Fig. S3|** Carbon flower size distribution after carbonization at 1000 °C **(a)**, 1250 °C **(b)**, and 1500 °C **(c)** and overview **(d)**. Error bars represent standard deviation over all particles.

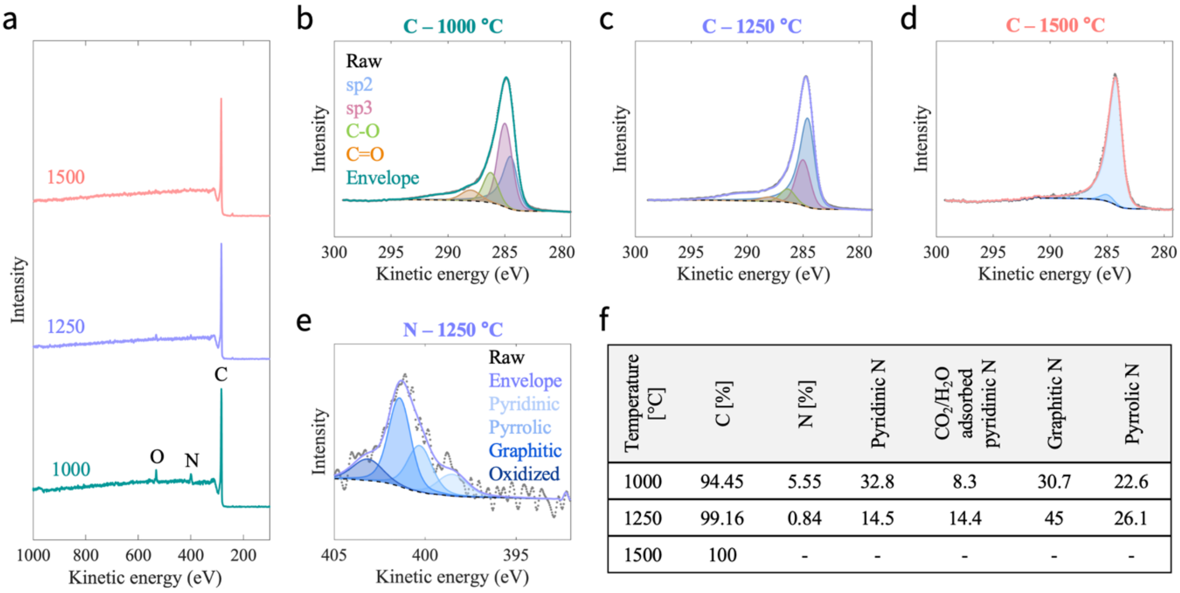


| Temperature [°C] | C [%] | N [%] | Pyridinic N | $CO_2/H_2O$ adsorbed pyridinic N | Graphitic N | Pyrrolic N |
|---|---|---|---|---|---|---|
| 1000 | 94.45 | 5.55 | 32.8 | 8.3 | 30.7 | 22.6 |
| 1250 | 99.16 | 0.84 | 14.5 | 14.4 | 45 | 26.1 |
| 1500 | 100 | - | - | - | - | - |

**Fig. S4| XPS spectra. (a)** Survey spectrum for CF carbonized at 1000, 1250, and 1500 °C. The expected C, N, and O peaks are indicated. High-resolution carbon scans on the **(b)** CF1000, **(c)** CF1250, and **(d)** CF1500. From left to right: C=O , C-O, sp3, sp2 carbon. **(e)** High-resolution nitrogen scans on the CF1250. From left to right: oxidized, graphitic, pyrrolic, pyrdinic nitrogen. **(f)** Table overview of composition.

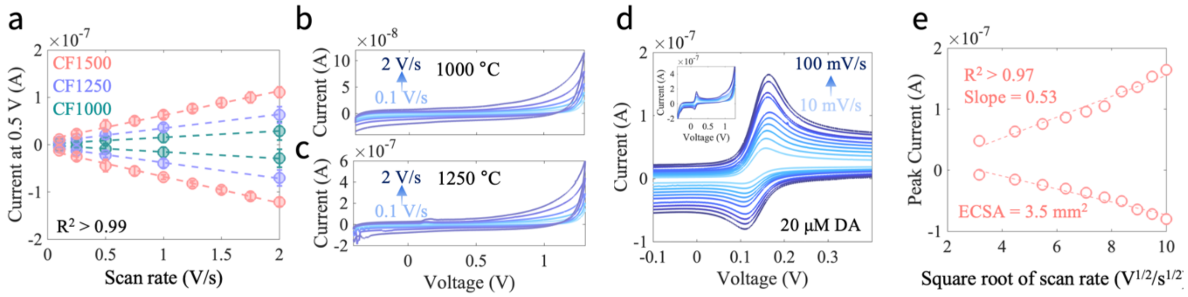


**Fig. S5| CV scans and ECSA. (a)** Capacitive current at 0.5V as a function of scan rate for CF1000, CF1250, and CF1500. Error bars represent 8 electrodes. Cyclic voltammograms of sensors fabricated from CF1000 **(b)** and CF1250 **(c)**. **(d)** CVs recorded with 20 µM dopamine for electrochemically active surface area (ECSA) estimation. The inset shows the full range. **(e)** Cathodic and anodic peak currents from (d).

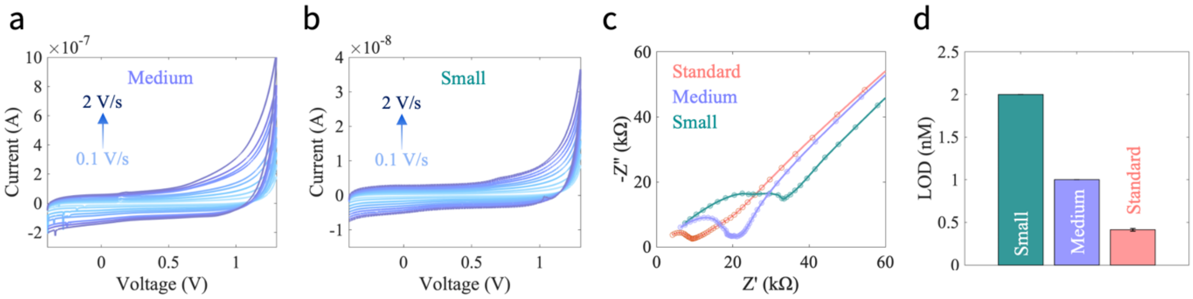


**Fig. S6| Effect of electrode size.** CV scans of medium (1.5 × 0.5 mm) **(a)** and small sensors (0.5 × 0.5 mm) **(b)**. **(c)** Impedance of all three sensors. **(d)** LOD for estradiol detection with different sizes.

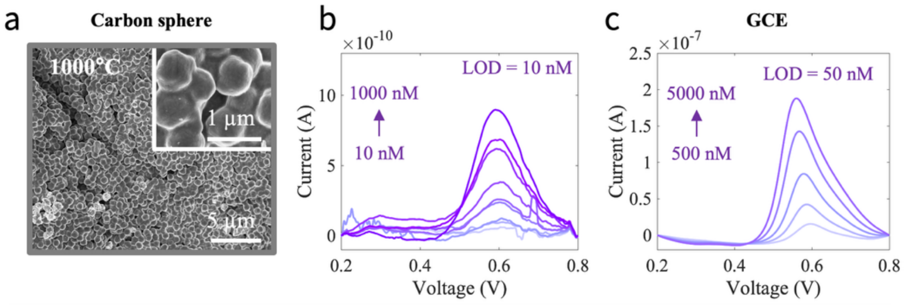


**Fig. S7| Carbon comparison.** SEM image of carbon spheres **(a)** and their estradiol detection **(b)**. **(c)** Estradiol detection with glassy carbon electrodes (GCE).

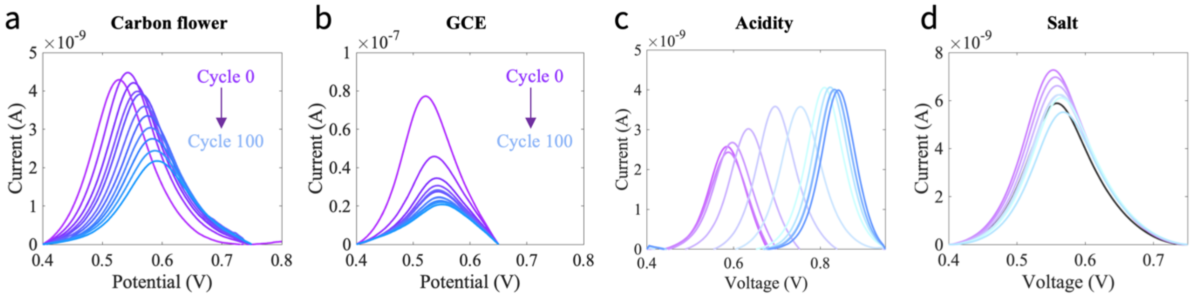


**Fig. S8| Robustness.** Raw data of repetitive cycling, showing every 10th cycle, for carbon flower electrodes **(a)** and GCE **(b)**. Raw data scans for various acidity **(c)** and salt **(d)** contents.

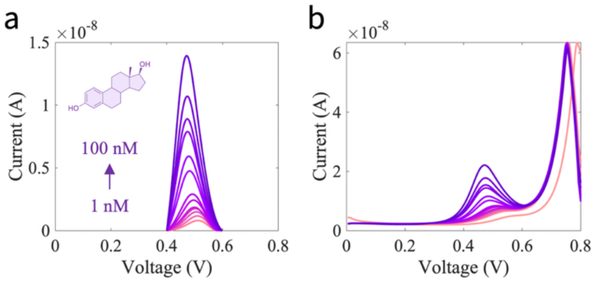


**Fig. S9| Artificial saliva.** Raw **(a)** and background subtracted **(b)** signal in artificial saliva.

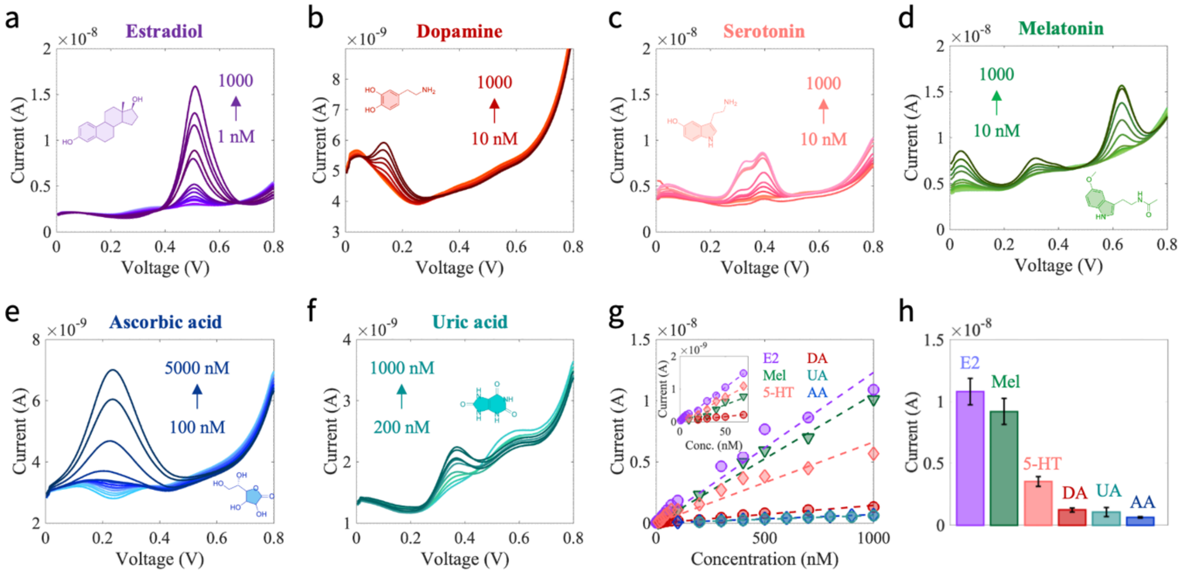


**Fig. S10| Raw data - detection of various biomarkers using carbon flower sensors.** Detection of estradiol (**a**), dopamine (**b**), serotonin (**c**), melatonin (**d**), ascorbic acid (**e**), and uric acid (**f**) for the indicated concentration ranges. **(g)** Linear correlation between peak current and concentration for each biomarker, presented for different concentration ranges. **(h)** Peak currents at 1000 nM for each analyte, averaged over eight channels.

**Table S1|** Carbon-based electrochemical sensors reported on Google Scholar since 2020 for detection of 5-HT, E2, DA, Mel, UA, and AA. Only sensors with limits of detection < 1 µM are included. Biosensors and molecularly imprinted polymer (MIP) sensors were excluded. Works are ordered by LOD. This work is indicated by *

| | Material Composition | LOD [nM] | Ref |
|---|---|---|---|
| **Dopamine** | 3DFG MEA Fuzzy Graphene | 0.36 | [36] |
| | GO-AgNPs / MWCNTs | 2.65 | [37] |
| | **Carbon flowers** | **4** | * |
| | $Fe_2O_3$ / LIG | 5.6 | [19] |
| | $Fe_2O_3$ / microelectrode | 8.76 | [38] |
| | Carbon cloth | 10 | [39] |
| | Fe/$Fe_3C$ / CNT | 15 | [40] |
| | AuNBP / MWCNT / GCE | 15 | [41] |
| | L-tryptophan / graphene /GCE | 60 | [42] |
| | rGO / Ppy | 61 | [43] |
| | Quantum Dot / MWCNT | 95 | [44] |
| | 3D hierarchical mesoporous carbon | 100 | [45] |
| | NSPAC | 100 | [46] |
| | rGO | 110 | [47] |
| **Estradiol** | **Carbon Flowers** | **0.3** | * |
| | $CeO_2$ NPs / graphite | 1.3 | [48] |
| | Graphene-coated silver nanoparticles / graphitic carbon nitride / GCE | 2 | [49] |
| | Cadmium molybdate / carbon nanospheres | 3 | [50] |
| | rGO-AuNPs / CNT / SPE | 3 | [51] |
| | α-$Fe_2O_3$-CNT / GCE | 4.4 | [52] |
| | Poly-L-Tyrosine / AuNCs / PDA-CNTs | 7.1 | [53] |
| | Wrinkled mesoporous carbon nanomaterials | 8.3 | [54] |
| | 3D graphene foam / SPE | 18 | [55] |
| | Au / Methylene blue / SPE | 41 | [56] |
| | Cobalt ferrite / rGO | 42 | [57] |
| | Polyaniline / carbon dot / GCE | 43 | [58] |
| | $Eu_2O_3$ / rGO | 50 | [59] |
| **Serotonin** | Graphitic Carbon Nitride / GCE | 0.15 | [60] |
| | **Carbon Flowers** | **1** | * |
| | Ultrafine $Fe_3O_4$ nanoparticles / carbon spheres | 4 | [61] |
| | $Ti_3C_2T_x$ / rGO | 10 | [62] |
| | $MnO_2$ / graphene | 10 | [63] |
| | FeC / AuNPs / MWCNT | 17 | [64] |
| | SPCE / MWCNT / Antimony Oxide NP | 24.6 | [65] |
| | Pd NPs / carbon dots / silica hybrid nanostructures | 33 | [66] |
| | $Nb_2CT_x$ / Protonated Carbon Nitride Nanocomposite | 63.24 | [67] |
| | Carbon nanohorns / GCE | 90 | [68] |
| | Nafion /CNT / CFME | 140 | [69] |
| | α-$MnO_2$ Nanorods / GCE | 140 | [70] |
| **Melatonin** | **Carbon Flowers** | **1** | * |
| | Carbon nanofiber arrays / FeCo alloys | 2.7 | [71] |
| | Porous flower-like hematite (α-$Fe_2O_3$) NPs / graphitic carbon nitride | 4 | [72] |
| | Porous Graphene / Au | 8.2 | [73] |
| | Sonogel-Carbon electrode material / Au NPs | 8.4 | [74] |
| | CNT microelectrodes | 10 | [75] |
| | poly (glycine) / carbon paste | 730 | [76] |

Abbreviations: GCE: glassy carbon electrode, LIG: laser induced graphene, CFME: carbon fiber microelectrodes, (MW)CNT: (multi-walled) carbon nanotubes, (r)GO: (reduced) graphene oxide, NP: nanoparticle, SP(C)E: screen printed (carbon) electrode, NSPAC: nitrogen- and sulfur-doped, porous, activated carbon.
